# When Does Molecular Dynamics Improve RNA Models? Insights from CASP15 and Practical Guidelines

**DOI:** 10.1101/2025.08.22.671717

**Authors:** Chandran Nithin, Smita P Pilla, Sebastian Kmiecik

## Abstract

Molecular dynamics (MD) simulations are increasingly applied to refine biomolecular models, yet their practical value in RNA structure prediction remains unclear. Here, we systematically benchmarked the effect of MD on RNA models submitted to the CASP15 community experiment, using Amber with the RNA-specific χOL3 force field. Across 61 models representing diverse targets, we find that short simulations (10–50 ns) can provide modest improvements for high-quality starting models, particularly by stabilizing stacking and non-canonical base pairs. In contrast, poorly predicted models rarely benefit and often deteriorate, regardless of their CASP difficulty class. Longer simulations (>50 ns) typically induced structural drift and reduced fidelity. Based on these findings, we provide practical guidelines for selecting suitable input models, defining optimal simulation lengths, and diagnosing early whether refinement is viable. Overall, MD works best for fine-tuning reliable RNA models and for quickly testing their stability, not as a universal corrective method.

## Introduction

RNA plays a central role in numerous cellular processes beyond its classical function as a messenger. It acts as a regulatory molecule, structural scaffold, and enzyme, and is increasingly recognized as a valuable therapeutic target (1–3). These diverse functions are tightly linked to RNA’s complex and dynamic three-dimensional (3D) structure, which is stabilized by intricate interactions such as noncanonical base pairing, base stacking, and long-range tertiary contacts (4–8). Accurately modeling RNA 3D structures remains a major challenge due to their intrinsic flexibility and the sparse availability of high-resolution experimental data (9, 10).

Recent progress in machine learning–based RNA structure prediction has significantly improved our ability to capture global folds (10–14). However, predicting local interactions and tertiary motifs with atomic-level accuracy remains difficult (11, 12, 15–17). As a result, predicted models often deviate from native conformations in ways that compromise their utility for mechanistic interpretation or structure-based design (9, 18, 19).

Among post-prediction refinement techniques, molecular dynamics (MD) simulations are perhaps the most widely used and intuitively appealing strategy. By modeling atomic-level fluctuations over time, MD can, in principle, relax strained geometries and reveal more native-like conformations. While MD has been successfully applied to protein refinement, its effectiveness for RNA model improvement remains unclear. All-atom MD simulations are computationally intensive, and their outcomes depend critically on the accuracy of the underlying force field (5, 20–22).

Drawing on our experience in RNA structure prediction assessments such as CASP (11–13) and RNA-Puzzles (15, 23), we observed that many modeling groups incorporate MD as a post-prediction refinement step. However, in most cases, the specifics of these simulations—particularly their timescales—are not clearly described, making it difficult to evaluate how MD actually contributes to model improvement.

Several studies have shown that short MD simulations—on the order of tens of nanoseconds—can help relax local geometric artifacts, correct base stacking, and reduce steric clashes in RNA models, particularly when starting from near-native conformations or structures derived from experimental restraints (24–27). However, the ability of longer, unbiased MD trajectories to systematically improve the global accuracy of RNA structural models remains unclear. In some cases, prolonged simulations may even lead to structural divergence, particularly when the starting model contains significant topological or base-pairing errors (5, 27). Despite these concerns, a common belief inherited from protein modeling persists—that longer simulations are inherently more effective at refining structure, due to greater conformational sampling. Whether this assumption holds true for RNA, given its distinct folding landscape and sensitivity to force-field limitations, has not yet been systematically addressed.

In this work, we use the AMBER ff99bsc0χOL3 (χOL3) force field (28–31), which is currently the best-supported and most extensively benchmarked parameter set for RNA molecular dynamics. Benchmark studies (32, 33), including those reviewed by Šponer et al.(5), have shown that χOL3 provides substantial improvements over earlier RNA force fields— particularly in modeling backbone torsions, canonical base pairs, and A-form geometries. χOL3 remains, to this day, the most widely validated and empirically supported force field for RNA simulations (20). While some limitations persist—such as the overstabilization of stacking interactions and limited accuracy for specific noncanonical motifs— χOL3 continues to offer the most balanced tradeoff between reliability and broad applicability for general RNA modeling.

To address the questions outlined above, we systematically investigate the effect of continuous, all-atom MD simulations—extending up to 300 nanoseconds—on RNA structure quality. We used a representative set of top-ranked RNA models from CASP15(11, 14), reflecting the current state of the art in blind RNA structure prediction. These models span a range of initial accuracies and CASP-defined difficulty categories, providing a realistic benchmark for assessing refinement methods. We assess whether MD can refine, stabilize, or degrade RNA structures over time, focusing especially on how structural quality evolves at different time points and whether early-stage dynamics (e.g., the first 50 ns) are predictive of long-term refinement outcomes. Our goal is to provide, to our knowledge, the first systematic CASP-scale evidence clarifying when MD is beneficial and when it is detrimental in RNA refinement, and to determine how simulation length modulates its utility.

## Materials and Methods

### Selection of CASP15 RNA Models for Analysis

We conducted MD simulations on all RNA targets from the CASP15 experiment, with the exception of cases involving RNA–protein complexes (R1189, R1190) or those too large for feasible simulation (R1138, 720 nt). This resulted in a total of 61 RNA models for nine targets: R1107, R1108, R1116, R1117, R1126, R1128, R1136, R1149, and R1156 (full list in Supplementary Table S1). For each target, we included the experimentally determined reference structure along with one model from each of five top-performing predictor groups (TS232, TS287, TS081, TS128, TS416) and a model submitted by our own group (TS392), yielding seven models per target, except for R1108 and R1116, which had six models available. This dataset spans a diverse range of sizes, folds, and prediction qualities, and includes representatives from all four CASP15 difficulty categories (11, 14) — Easy (well-structured RNAs with accurate predictions, typically supported by homologous structures in the PDB), Medium (partially correct global folds, where structural similarity could be inferred from related functions), Difficult (large or topologically complex RNAs lacking homologous structures), and *Non-natural* (synthetic designs with novel folds) — while ensuring direct comparability to native structures (see Supplementary Table S1).

### Molecular Dynamics Simulations

MD simulations were performed using the Amber 22 package (34), with the AMBER ff99bsc0χOL3 (χOL3) RNA force field. This force field was chosen based on its extensive validation and widespread use in RNA modeling; see the Introduction for rationale and references regarding force field performance.

A 2 fs integration timestep was used for all simulation steps. Bonds involving hydrogen atoms were constrained using SHAKE (ntc=2, ntf=2), with SETTLE applied for rigid water molecules (35, 36). A non-bonded cutoff of 12 Å was set. Long-range electrostatic interactions were calculated using the Particle-Mesh Ewald (PME) method (37). Energy minimization and all subsequent simulation steps were facilitated using the CUDA-accelerated PMEMD (38–40).

Initial structures were prepared using tleap (41). Each system was enclosed in a truncated octahedral box with a 10 Å buffer. The system was neutralized with Na+ ions. Solvation was performed using the TIP3P water model (42).

Energy minimization was conducted in two stages of 10,000 cycles each. In the first stage, positional restraints were applied to the backbone phosphorus (P) and oxygen atoms (OP1, OP2) of the RNA backbone with a restraint weight (restraint_wt) of 20.0 kcal mol^−1^ Å^−2^. The second stage was performed without any restraints.

The system was gradually heated from 100 K to 300 K over 500 ps (250,000 steps). During heating, positional restraints with a weight of 20.0 kcal mol^−1^ Å^−2^ were maintained on the RNA backbone atoms. The heating process was controlled using the Langevin thermostat (43) with a collision frequency (gamma_ln) of 5.0 ps^−1^, and random seed (ig=-1) for stochastic dynamics. The temperature was ramped linearly from 100 K to 300 K during this simulation.

Following heating, the system underwent a four-phase equilibration process under NVT conditions. Initially, the system was equilibrated for 200 ps with positional restraints applied to the backbone atoms (O, OP1, OP2, P) using a restraint weight of 10.0 kcal mol^−1^ Å^−2^. The temperature was maintained at 300 K using the Langevin thermostat (gamma_ln=5.0). This was followed by an additional 200 ps equilibration with the restraint weight reduced to 5.0 kcal mol^−1^ Å^−2^, and another 200 ps with the restraint weight further lowered to 1.0 kcal mol^−1^ Å^−2^. Finally, a 2 ns equilibration was conducted without restraints.

The production phase of the simulation was conducted under constant pressure conditions using the NPT ensemble. A total of 300 ns of simulation time was performed with a 2-fs time step, resulting in 150 million steps (nstlim=150000000). The system was maintained at 298 K (temp0=298.0) using the Langevin thermostat (ntt=3) with a collision frequency (gamma_ln) of 1.0 ps−1. The pressure was set to 1.0 atm (pres0=1.0) with isotropic position scaling (ntp=1) and a pressure relaxation time (taup) of 2.0 ps. The simulation was set to restart from previous coordinates and velocities (irest=1) with random seed initialization (ig=-1).

To characterize and compare the conformational free energy landscapes, a multi-step analysis protocol was applied to the simulation data for each RNA target. First, all solvent and ion coordinates were removed from the MD trajectories, and the data for all simulated models and the experimental reference for a given target were pooled into a single combined ensemble. The internal structural dynamics of this ensemble were described using a high-dimensional feature set consisting of the pairwise distances between the centers of mass of the heavy atoms of every nucleotide. This feature data was then subjected to a two-step dimensionality reduction protocol. The feature sets were first projected onto their top 500 principal components using Principal Component Analysis (PCA) to denoise the data, followed by time-lagged independent component analysis (tICA). A lag time of 20 ns was chosen for tICA to identify the slowest, most functionally relevant collective motions, and the top 10 independent components (tICs) were retained for analysis. The final two-dimensional free energy landscapes were constructed by projecting the combined-ensemble trajectories onto pairs of these tICs, and the Potential of Mean Force (PMF) was calculated via Boltzmann inversion. While the landscapes projected onto the first two components (tIC1 and tIC2) capture the most dominant large-scale motions (Supplementary Figures S4, S5, S7A, S8A), the projections onto tIC3 and tIC4 provided superior visual separation of the key states, most clearly distinguishing the native ensemble from the distinct non-native basins sampled by the models (Figures 4A, 5A, Supplementary Figures S7B, S8B).

### Evaluation Metrics

To quantify structural similarity between predicted RNA models and experimental reference structures, we used root-mean-square deviation (RMSD) and Interaction Network Fidelity (INF). RMSD was computed for all heavy atoms after least-squares superposition of the models onto the native structures using PyMOL (Schrödinger, Inc.) (44). When multiple biological assemblies were available, RMSD was evaluated against all reference assemblies. INF was calculated with *rna-tools* (45) based on ClaRNA annotations (46) and measures the extent to which predicted base-pairing and stacking interactions reproduce those of the reference (47). The metric is derived from the Matthews correlation coefficient, with INF = 1 indicating perfect agreement and INF = 0 indicating no overlap. We report the overall INF score (INF_all) as well as interaction-specific components: INF_stack for stacking, INF_wc for canonical Watson–Crick pairs, and INF_nwc for non-Watson–Crick pairs.

## Results and Discussion

### Global Structural Stability and the Limits of Refinement

To assess the potential of MD simulations for refining the global fold of RNA models, we monitored the evolution of the root-mean-square deviation (RMSD) relative to the corresponding experimental structures (Figure 1). RMSD, calculated over all non-hydrogen atoms, provides a direct measure of the overall similarity between a model and its native fold. Figure 1A illustrates the diversity of behaviors across individual trajectories, whereas Figure 1B shows the smoothed median RMSD for each target, highlighting the average structural trend.

**Figure 1.**
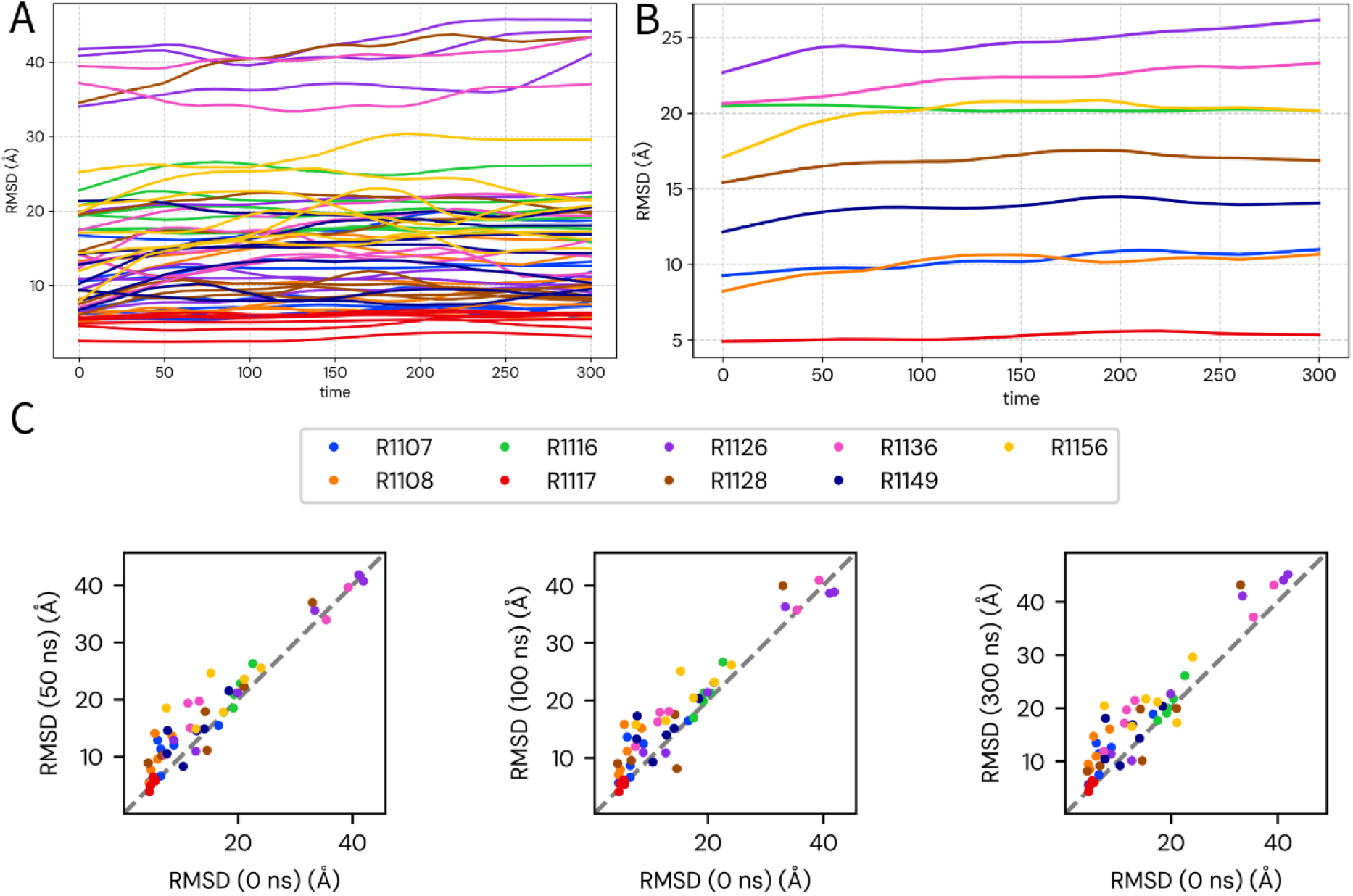
Global structural stability of models assessed by RMSD. Time series plots and scatter plots of the root-mean-square deviation (RMSD, in Å) for the 52 submitted models relative to their respective experimental native structures. (A) RMSD trajectories for individual models; each thin colored line represents a single simulation, color-coded by RNA target, illustrating the distribution of model quality and its temporal evolution. (B) Median RMSD trajectories for each target; each thick colored line represents the median RMSD of 5–6 models for that target. (C) Scatter plots comparing RMSD at three time points (50, 100, and 300 ns) with the initial RMSD (0 ns). The dashed grey diagonal represents no change; points above the line indicate higher RMSD relative to the starting model.

A consistent observation across our dataset is the lack of marked global structural refinement during the course of the simulations. As shown by the median trajectories in Figure 1B, none of the targets display a substantial decrease in RMSD relative to the native structure. Instead, most models remain approximately stable, maintaining a relatively flat RMSD profile, or show a gradual increase in RMSD suggestive of structural drift. This tendency is seen even for the highest-quality initial predictions: the individual trajectories in Figure 1A illustrate that models starting with the lowest RMSD values for their respective targets did not converge more closely toward the native structure. For example, the best models for the “Easy” target R1117 (red lines) begin at ∼5 Å RMSD and fluctuate around this value for the entire 300 ns, showing stability but no clear improvement.

This trend is also apparent in the scatter plots (Figure 1C). Across all three time points, most models lie either on or above the diagonal, indicating stability or gradual drift away from the native structure. The stability of the best predicted models for targets such as R1117 (red points) and the designed RNA R1128 (brown points) is reflected by their tight clustering along the diagonal. By contrast, models for more challenging targets like R1116 (green points) show a progressive upward deviation from the line, particularly between 50 ns and 300 ns, consistent with cumulative structural divergence.

In summary, the RMSD analysis suggests that, under the conditions tested here, unbiased MD simulations generally did not bring models closer to the native global fold. Local fluctuations were observed, but the large-scale corrective motions needed for global refinement were rare. In most cases, model quality was preserved, while some trajectories showed gradual divergence. These results highlight that, with the current protocol and force field, the main value of MD lies more in stability assessment than in global improvement. Because RMSD reflects only overall geometric similarity and is insensitive to detailed base-pairing patterns, we next turned to Interaction Network Fidelity (INF) to evaluate whether local RNA interactions were preserved or improved during the simulations.

### Global Trends in Interaction Network Fidelity (INF) Across MD Simulations

INF analysis provided a complementary view of RNA model behavior, revealing interaction-specific trends that were not apparent from RMSD alone (Figure 2). Canonical Watson–Crick base pairs (INF_wc) were highly stable across nearly all targets, with median scores consistently above 0.95 throughout the 300 ns trajectories and narrow interquartile ranges indicating strong model consensus. Stacking interactions (INF_stack) were similarly robust, with only minor declines observed in most cases (Supplementary Figures S1, S2).

**Figure 2:**
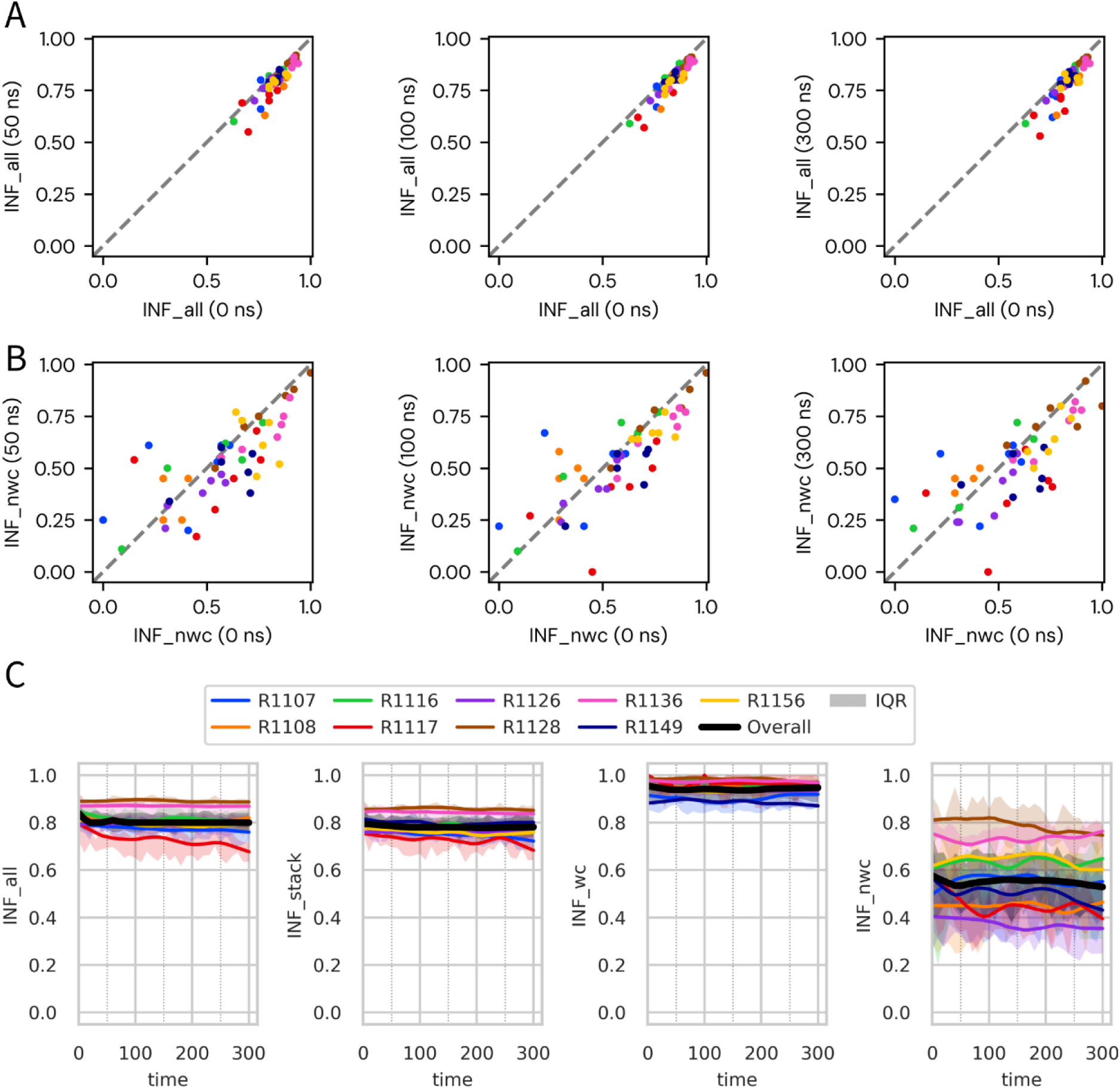
Dynamics of RNA interaction networks assessed by Interaction Network Fidelity (INF). (A) Scatter plots comparing overall interaction fidelity (INF_all) at 50, 100, and 300 ns against initial values (0 ns). Each point represents a single model, color-coded by RNA target; the dashed diagonal indicates no change, with points below the line reflecting degradation. (B) Equivalent scatter plots for non-canonical interactions (INF_nwc), highlighting both gains (above the line) and losses (below the line). (C) Time series of median INF scores for all four components: overall (INF_all), stacking (INF_stack), Watson–Crick (INF_wc), and non-Watson–Crick (INF_nwc). Each colored line represents the LOWESS-smoothed median across all predicted models for a given target (excluding the experimental structure), and shaded areas indicate the interquartile range. Stacking and Watson–Crick components are shown in detail in Supplementary Figure S2.

In contrast, non-canonical interactions (INF_nwc) were substantially more dynamic and constituted the main source of variability in overall fidelity (INF_all). Depending on the target, INF_nwc either improved or deteriorated, leading to corresponding trends in INF_all. For example, the “Medium” target R1107 and the “Difficult” targets R1116 and R1156 showed upward trajectories in INF_nwc, reflecting partial recovery of initially weak non-canonical networks. In R1107, this effect was driven largely by model TS416, which substantially repaired its non-canonical contacts, whereas other models showed smaller gains or modest declines. A similar pattern was evident in R1116 (Supplementary Figure S1), where improvement in one model offset losses in others, and in R1156, which displayed gradual increases from a low starting point.

By contrast, other targets exhibited widespread loss of non-canonical interactions. The “Easy” target R1117 showed consistent declines across nearly all models, including those with initially strong networks. The designed RNA R1126 displayed a similar but less pronounced pattern, whereas R1128 remained stable, with high INF_nwc scores maintained throughout the simulations.

Overall, these findings show that non-canonical interactions are the most variable aspect of RNA model fidelity during MD. Depending on the starting accuracy and simulation conditions, they can either support refinement by recovering native contacts or undermine fidelity through progressive loss of interactions.

### Impact of Starting Structure Quality on Interaction Network Fidelity (INF) During MD Simulations

To examine how starting accuracy influenced model behavior during simulation, we used the initial RMSD to the native structure as a global measure of fold quality. The highest-quality group (initial RMSD <5 Å) included all models for the “Easy” target R1117 (2.0–4.7 Å), top models for the “Medium” targets R1107 and R1108 (e.g., TS232 at 4.5 Å), and the designed target R1128 (TS232 at 4.3 Å). Despite their close overall agreement with experimental structures, these models displayed contrasting outcomes at the level of local interactions. R1117 models showed a rapid loss of non-canonical contacts (INF_nwc), whereas R1128 and R1108 models maintained robust interaction networks throughout the trajectories (Supplementary Figures S1). Thus, an excellent initial RMSD was not a sufficient predictor of stability; rather, the energetic plausibility of the non-canonical network emerged as the critical determinant.

Models with initial RMSD values between 5 and 10 Å represented a mixed tier, spanning “Medium,” “Difficult,” and “Non-natural” targets. Outcomes were highly variable: for example, R1156 models in this range (e.g., TS128 at 5.4 Å) frequently showed refinement of their non-canonical networks, whereas other models with comparable global folds displayed pronounced volatility or degradation in INF_nwc scores.

The 10–20 Å RMSD range, which included most models for the “Difficult” targets R1116 (e.g., TS081 at 12.7 Å) and R1156 (e.g., TS081 at 17.1 Å), was the regime most conducive to refinement. These models often began with substantial errors in non-canonical contacts but were sufficiently close to the native state for MD to promote rearrangement of tertiary interactions. As a result, they were the group most frequently showing improvement in INF_nwc (Supplementary Figure S1), suggesting that intermediate-accuracy models possess both the flexibility and proximity to native geometry needed for force-field–driven corrections.

By contrast, models starting more than 20 Å from the native structure were generally too far to benefit from refinement. This category included models with major topological errors, such as R1116 (TS392 at 21.2 Å) and the designed targets R1126 and R1136. Their interaction networks remained in poor, non-native states, typically starting with low INF_nwc scores and showing little change. These structures were effectively “stably wrong,” neither improving substantially nor undergoing catastrophic collapse.

### Improvements and Degradation of INF the Early Phase of Simulation

A consistent temporal pattern across simulations is that the most substantial changes in interaction networks—whether beneficial or detrimental—occur within an early window, typically the first 50 ns (Figure 3). This phase reflects the immediate response of the models to the force field and is particularly pronounced in the non-canonical interaction network.

**Figure 3:**
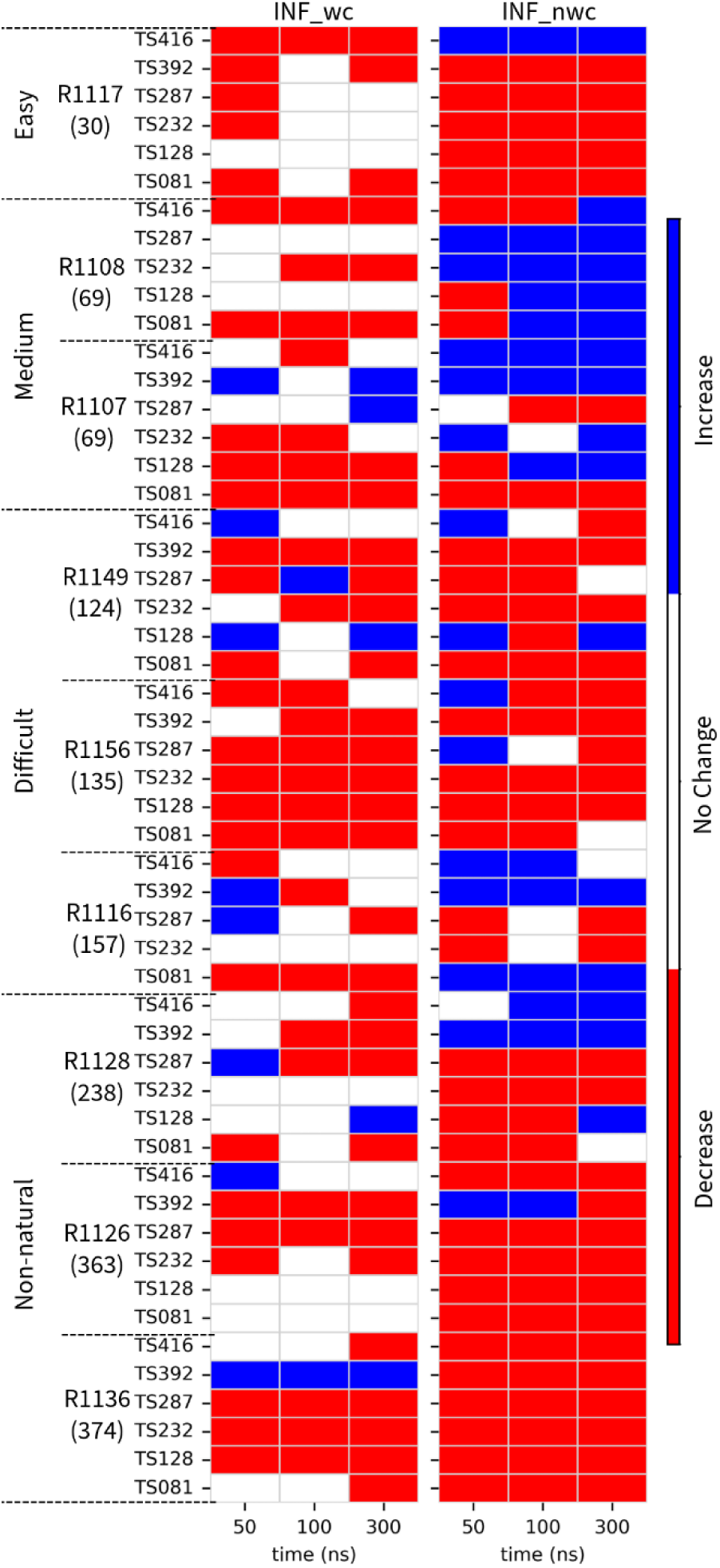
Early-phase changes in interaction networks. Heatmap showing changes in INF scores (ΔINF) for Watson–Crick (INF_wc, right panel) and non-Watson–Crick (INF_nwc) interactions at 50, 100, and 300 ns relative to the starting structure. Blue cells indicate improvement (ΔINF > 0), red cells indicate loss (ΔINF < 0), and white cells indicate no change. The contrast between the two components is evident: Watson–Crick contacts remain largely stable, while non-canonical interactions are highly dynamic, exhibiting both refinement and degradation.

**Figure 4:**
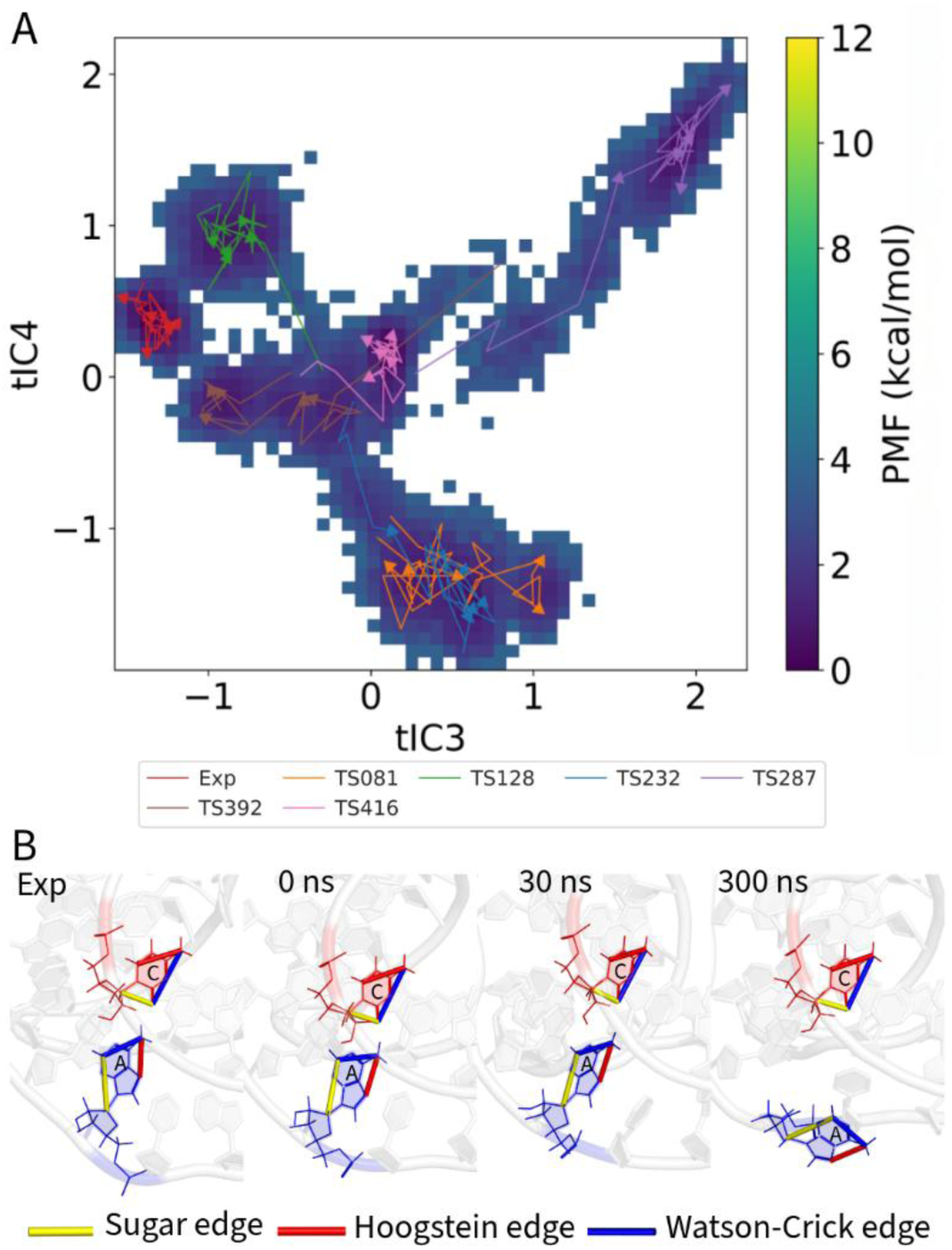
Structural dynamics of the “Easy” target R1117. (A) Combined two-dimensional potential of mean force (PMF) landscape for all MD simulations of the PreQ1 class I type III riboswitch (R1117), projected onto the third (tIC3) and fourth (tIC4) time-lagged independent components. Free energy is shown in kcal/mol (purple = low, yellow = high). Arrows trace conformational trajectories of each model from 0 to 300 ns; the experimental reference is shown in red, and CASP models are color-coded by target. (B) Example of non-canonical base-pair dynamics in model TS416. Snapshots illustrate the cis sugar-edge/Watson–Crick (SW) A– C interaction at four stages: experimental reference, starting model (0 ns), and MD snapshots at 30 ns and 300 ns. Edges are colored according to the Leontis–Westhof scheme: sugar (yellow), Hoogsteen (red), Watson–Crick (blue).

For the “Easy” target R1117, non-canonical fidelity (INF_nwc) illustrates a transient correction followed by rapid decline. Median scores improved sharply during the first 10 ns but quickly decayed, with most gains lost by 50 ns (Supplementary Figures S1). This behavior suggests that fast, local adjustments to tertiary contacts can occur, but these are often not energetically stable enough to persist.

Conversely, the early window is also where lasting improvements take root. For the “Difficult” targets R1116 and R1156, which exhibited overall gains in INF_nwc, a large fraction of the improvement was established within the first 50–100 ns (Supplementary Figure S1). Similarly, the “Medium” target R1107 achieved its modest increases in INF_nwc largely during this initial phase (Supplementary Figure S1).

The same early period also encompasses the most severe degradations. For instance, models TS392 for R1117 and R1116 experienced precipitous declines in INF_nwc to near-zero within 10 ns, while others, such as the designed target R1126, showed a steady downward trajectory already evident by 50 ns (Supplementary Figure S1).

By contrast, models that began in particularly stable states, such as those for the designed target R1128, showed minimal early-phase change. Their INF_nwc values remained consistently high from the outset, indicating that they lacked the strained local contacts that drive rapid adjustment in other systems (Supplementary Figure S1).

Taken together, these observations indicate that early-phase changes largely reflect the rapid relaxation of high-energy strains present in the starting models. Once this initial relaxation is complete, trajectories enter a slower regime in which crossing larger energy barriers becomes limiting. This has practical implications: refinement strategies may benefit more from identifying and stabilizing short-lived improvements arising in the first tens of nanoseconds than from simply extending simulations over longer timescales.

### Case Studies of Specific Targets

To illustrate how the general trends observed in RMSD and INF analyses manifest at the level of individual systems, we present two representative case studies in the main text. These examples highlight contrasting outcomes: one in which initially accurate models undergo rapid loss of non-canonical interactions, and another in which imperfect starting structures achieve sustained refinement. Additional case studies covering the remaining CASP15 targets are provided in the Supplementary Information, where they further support the diversity of behaviors summarized here. By combining global analyses with illustrative examples, we aim to show how initial accuracy, interaction network composition, and early-phase responses collectively determine the trajectory of refinement during MD simulations.

### Easy target prone to rapid non-canonical collapse despite high starting quality

Target R1117 (PreQ1 class I type III riboswitch), classified as “Easy” in CASP15, began with several models of high starting accuracy (RMSD ≈ 2–5 Å)(11). Despite this favorable starting point, most models experienced rapid decline of non-canonical contacts within the first tens of nanoseconds. Projection of all trajectories onto a common two-dimensional potential of mean force (PMF) landscape using time-lagged independent component analysis (tICA) illustrates this divergence (Figure 4A). While the experimental reference structure (red arrow) remained confined to a single stable basin, predicted models explored multiple alternative states, with arrows tracing divergent paths away from the native minimum.

Global fidelity scores reinforce this picture. Watson–Crick base pairs (INF_wc) and stacking interactions (INF_stack) remained generally stable, but non-canonical fidelity (INF_nwc) declined sharply, driving the overall drop in INF_all (Supplementary Figure S1). Individual trajectories highlight this vulnerability: TS287 and TS128 both started with strong INF_all values (>0.8) yet lost non-canonical interactions over time, TS081 showed only transient improvement before degradation, and TS392 collapsed almost immediately with INF_nwc approaching zero. Even TS232, among the best starting models, displayed steady erosion of its non-canonical network.

Model TS416 (pink) provides a mechanistic illustration of these dynamics. It began with good Watson–Crick fidelity but very poor non-canonical fidelity (INF_nwc = 0.15). Within the first nanoseconds, a key sugar-edge/Watson–Crick 24A–16C base pair transiently reformed, briefly raising INF_nwc to 0.45 (Figure 4B). This improvement, however, was not sustained: by 50 ns the interaction had dissociated, accompanied by a decline in INF_all and weakening of stacking interactions.

Together, these examples demonstrate that even initially accurate models of R1117 are prone to early and often irreversible erosion of their non-canonical networks. Short-lived corrections can occur, but they are generally insufficient to maintain global fidelity. This case underscores the pivotal role of non-canonical interactions in determining whether refinement is achievable or whether models instead drift toward less native-like states during simulation.

### Medium target with rare but stable recovery of tertiary contacts

Target R1107, a ‘Medium’ difficulty human CPEB3 HDV-like ribozyme, included mostly stable models during MD, but only one (TS416) showed a rare and sustained improvement in non-canonical interactions that persisted throughout the 300 ns trajectory. CASP15 evaluators classified it as ‘Medium’ difficulty, with the best predicted models reaching RMSDs around 4.5 Å (11). To assess conformational changes during simulation, we projected each model’s trajectory onto a combined 2D potential of mean force (PMF) landscape (Figure 5A, Supplementary Figure S4). The landscape reveals several distinct energy basins, with the experimental structure (red arrow) remaining in a well-defined native state. In contrast, CASP models followed diverse paths, showing variable stability and refinement outcomes.

**Figure 5:**
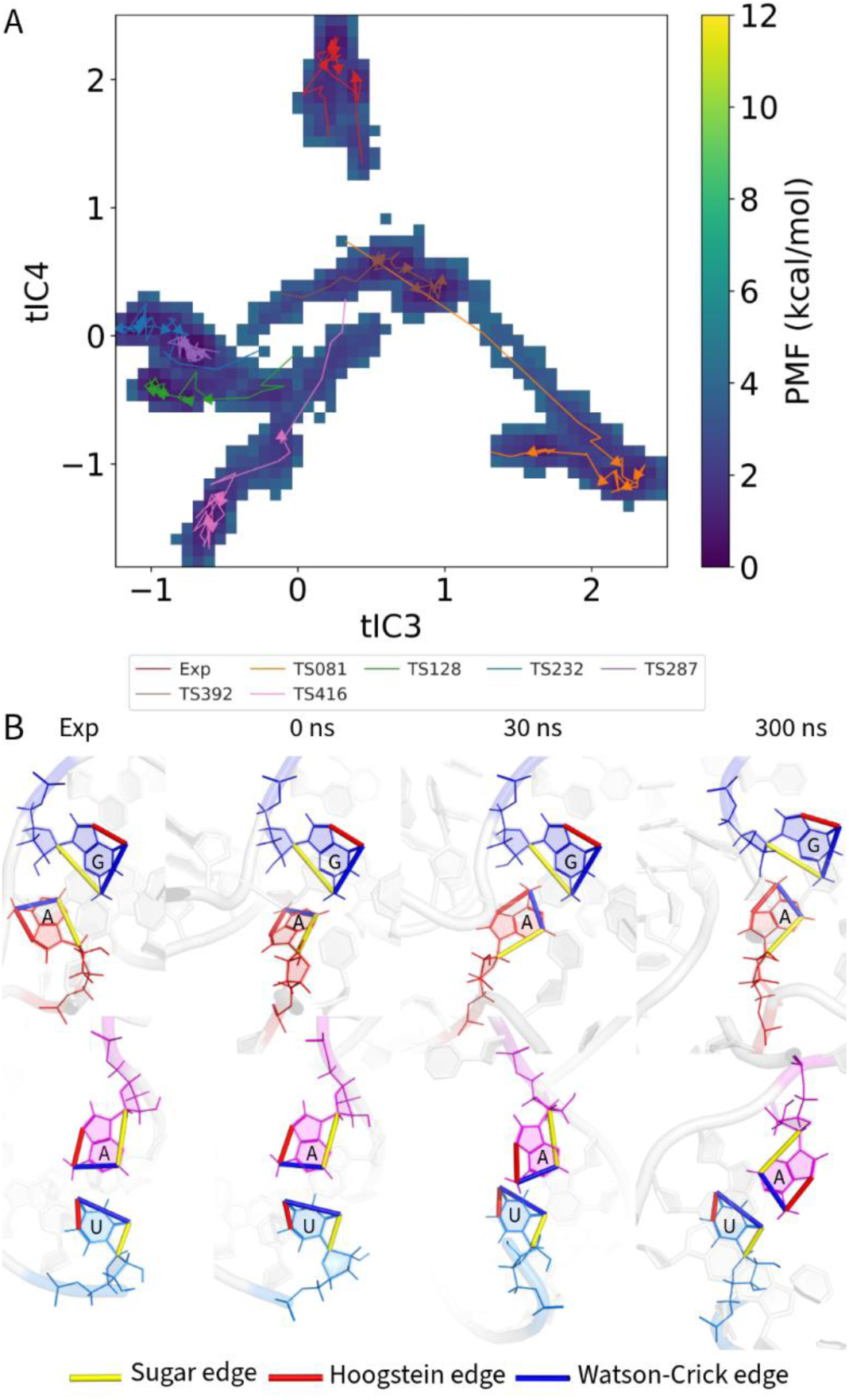
Selective refinement and general stability in the “Medium” target R1107. (A) Combined 2D potential of mean force (PMF) landscape for all MD simulations of the human CPEB3 HDV-like ribozyme (R1107), projected onto the third (tIC3) and fourth (tIC4) time-lagged independent components. Free energy is shown in kcal/mol, with low-energy basins in purple and high-energy regions in yellow. Arrows trace conformational trajectories of the experimental structure (red) and predicted CASP models. **(B)** Representative conformations from model TS392. Snapshots highlight two base pairs during simulation. Top: A28–A62, a native non-Watson–Crick pair, absent at 0 ns, recovered at 30 ns, and partially distorted at 300 ns. Bottom: A10–U69, a canonical Watson–Crick pair in the experimental structure and at 0 ns, which progressively deviated and was reclassified as non-native. Base edges are colored according to the Leontis–Westhof scheme: sugar (yellow), Hoogsteen (red), Watson–Crick (blue).

Model TS392 (orange) illustrates how apparent improvements in INF_nwc can mask countervailing effects (Figure 5B). One missing non-canonical pair, G28–A62 (sugar–Watson), reappeared by 30 ns and persisted through the trajectory, driving an initial rise in INF_nwc. At the same time, however, the canonical A10–U69 Watson–Crick contact progressively lost its native geometry, being reclassified as sugar–Hoogsteen at 50 ns and Watson–Watson trans at 300 ns. By the end of the run, an additional non-native C6–C32 (sugar–Hoogsteen) pair had emerged, further penalizing fidelity through false positives. Thus, although INF_nwc increased relative to the starting model, the gain came from a delicate balance of recovering one native tertiary contact while simultaneously distorting or inventing others.

Among the six predictions, TS416 stood out. It began with modest global quality (INF_all = 0.76) and a poor non-canonical network (INF_nwc = 0.22), but during simulation INF_nwc nearly tripled to 0.61, representing a substantial recovery of tertiary contacts. Although the overall INF_all rose only slightly to 0.78, this trajectory demonstrates that meaningful refinement is possible under favorable conditions. In contrast, the remaining models showed no comparable improvement. Some, such as TS081, deteriorated steadily, while others (TS232, TS287, TS128) began from relatively high fidelity and exhibited only minor declines. These small drifts are reflected in the narrow interquartile range in Supplementary Figure S1, emphasizing that stability was common but true refinement was exceptional.

Taken together, R1107 illustrates that while MD can occasionally enable sustained repair of non-canonical interactions, as in TS416, such events are rare. For most medium-quality models, prolonged MD was neutral at best and often led to gradual erosion of interaction fidelity.

### Implications for RNA Structure Modeling and Refinement

The contrasting behaviors of R1117 and R1107 highlight a central lesson of this study: the fate of an RNA model during MD is dictated more by its initial interaction network than by its CASP-assigned difficulty category. Even ‘Easy’ targets with excellent global folds, such as R1117, can undergo rapid and irreversible collapse of non-canonical contacts, while models of intermediate quality, such as R1107 TS416, may achieve partial and durable recovery of tertiary interactions. This underscores that starting quality—not category—is the primary predictor of MD outcomes. While force field artifacts may contribute to some observed degradation patterns, the systematic nature of our findings across diverse targets suggests that the limitations primarily reflect the inherent challenges of RNA refinement rather than specific χOL3 parametrization issues.

These findings have several practical implications. First, RMSD alone is not a reliable indicator of stability. Models with low RMSD but incorrect non-canonical geometry often deteriorated quickly, whereas some with higher RMSD but flexible tertiary contacts showed refinement potential. Second, the early phase of simulation (≤50 ns) is decisive: improvements, when they occur, emerge rapidly, while later stages mostly preserve the status quo or allow slow erosion. Thus, short simulations can act as an efficient filter to distinguish stable from unstable models. Finally, MD should be viewed less as a general refinement engine and more as a diagnostic tool: it is well-suited for confirming the robustness of strong predictions or revealing hidden fragilities, but it rarely transforms poor models into native-like structures.

In practice, this means that RNA modeling pipelines should emphasize generating the highest quality starting structures possible, then applying short MD simulations selectively as a stability screen. Prolonged unrestrained dynamics, by contrast, often provide limited benefits and may exacerbate deviations. Future progress will likely depend on hybrid strategies— combining physics-based MD with targeted restraints, enhanced sampling, or integration of experimental data—to stabilize correct non-canonical geometries and prevent collapse. These findings may also inform the development of improved scoring functions for RNA models that incorporate short-term MD stability as a structural quality indicator, complementing traditional geometric and energetic assessments.

### Computational Cost vs. Structural Improvement

MD simulations are a powerful tool for exploring conformational landscapes, but their application for refining RNA structures requires careful consideration of the significant computational cost versus the potential for improvement. All simulations were performed on state-of-the-art NVIDIA GH200 GPUs (96 GB), whose performance for MD is comparable to or exceeds that of previous high-end accelerators such as the A100 and H100, enabling efficient execution of long RNA trajectories. Despite this cutting-edge hardware, the full benchmark still required a total of 158.6 GPU-days to simulate 61 models across nine targets (Supplementary Figure S6), underscoring the heavy computational burden. While MD can successfully refine certain models, it is not a universal solution, and its efficacy is almost entirely dictated by the stability of the non-canonical interaction network.

The quality of the starting structure is a critical but not solitary determinant of the simulation outcome. A prime example of this complexity is the’Easy’ target R1117. Despite the high initial quality of many of its models, they experienced a near-universal and rapid degradation of their non-canonical networks, causing a drop in overall fidelity (Figure 1B). This indicates that even a well-predicted secondary structure is insufficient if the tertiary contacts are energetically strained. In stark contrast, two’Difficult’ targets, R1116 and R1156, represent the clearest cases of successful refinement. For these targets, models that started with very poor non-canonical networks (median INF_nwc ≈ 0.30) showed a steady and significant improvement over the 300 ns trajectory (Supplementary Figure S1). This demonstrates that MD can be a powerful corrective tool, but only when the target’s energy landscape permits relaxation toward the native state. Finally, for a well-predicted and architecturally regular target like the’Non-natural’ R1128, models with high initial fidelity remained exceptionally stable, showing that MD can also serve as a robust validation tool (Supplementary Figure S1).

The trade-off between computational expense and structural improvement becomes particularly acute for large RNA systems. As shown in Supplementary Figure S6, the simulation cost scales dramatically with RNA length: a single 300 ns trajectory for the smallest target (R1117, 30 nt) took less than a day, while one for the largest (R1136, 374 nt) required over six days. The total computational time for the R1136 ensemble exceeded 43 GPU-days, yet this investment only resulted in the preservation of its already moderately accurate non-canonical network, not a significant refinement (Supplementary Figures S1, S6). This highlights that for large RNAs, the high cost may not be justified by the expected improvement.

Considering these factors, our findings advocate for a strategic and diagnostic, rather than universal, application of MD for RNA model refinement. The stability of the non-canonical interaction network (INF_nwc) emerges as the most critical metric to monitor. For models that are already high quality, short simulations may be sufficient to validate stability. For flawed models, the INF_nwc trajectory can diagnose whether the structure is on a productive refinement pathway (as in R1116) or is trapped in a non-native state. Given the substantial computational cost, particularly for large ensembles and long RNAs, standard MD is best leveraged as a diagnostic tool to assess model robustness and identify regions amenable to targeted correction. Future refinements in modeling workflows may benefit from hybrid approaches—such as incorporating secondary structure restraints, using sparse experimental data, or refining only selected regions—so that computational effort is invested where it is most likely to yield structural improvement.

### Practical Guidelines for the Use of MD in RNA Structure Modeling

The results of this study allow us to propose practical guidelines for when and how MD simulations should be applied in RNA structure modeling workflows. The critical insight is that MD is not a universal refinement tool, but rather a diagnostic and selective strategy whose utility depends strongly on the initial quality of the model and its early trajectory during simulation. To support decision-making, we provide a set of general recommendations (Table 1) and a user-facing diagnostic checklist (Table 2).

**Table 1:** General guidelines for applying MD simulations depending on starting model quality.

| Starting model quality (relative to t=0 ns) | Expected MD behavior | Recommended strategy |
| --- | --- | --- |
| **High quality** (low RMSD, high INF_all and INF_nwc; stable INF_wc $\approx 1.0$) | Interaction network remains close to initial state; minor local relaxation possible. | Short MD (10–50 ns) sufficient for validation or small-scale optimization. No benefit from long runs. |
| **Intermediate quality** (moderate RMSD, partial INF_nwc loss, but not collapsed) | Early phase ($\leq 50$ ns) is diagnostic. Upward INF_nwc trend indicates refinement potential; downward trend predicts failure. | Run short MD first; extend only if early INF_nwc improves relative to t=0 ns. Terminate if decline is observed. |
| **Poor quality** (high RMSD, collapsed INF_nwc, large deviations from secondary structure) | Early catastrophic collapse (RMSD spikes, INF_wc loss, INF_nwc decline) or persistent instability. | Do not invest in MD. Terminate immediately. Use alternative modeling or restraints instead. |

**Table 2.** Practical checklist for assessing whether an RNA model is suitable for MD refinement (without a native reference).

| Diagnostic criterion | High-quality indicator (MD justified) | Low-quality indicator (MD not justified) |
| --- | --- | --- |
| Prediction consensus | Consistent across multiple algorithms / prediction methods | Divergent, inconsistent predictions |
| Secondary structure agreement | Matches reliable predictions or probing data (e.g., SHAPE, covariation) | Major conflicts with predicted or known secondary structure |
| Experimental restraints | Fits cryo-EM density, NMR NOEs, or other restraints | Poor fit or clear violations of restraints |
| Early RMSD trajectory (0–50 ns, relative to t=0) | Flat or modest drift | Sharp increase (>2–3 Å within tens of ns) |
| Early INF_wc (relative to t=0) | Remains $\approx 1.0$ (canonical base pairs stable) | Early decline (loss of canonical pairs) |
| Early INF_nwc (relative to t=0) | Flat or upward trend (recovery of tertiary contacts) | Early downward trend or collapse |
| Stacking interactions | Stable within first tens of ns | Rapid or progressive loss |

In practice, these guidelines emphasize the diagnostic value of the early phase of MD simulations. High-quality models, if stable within the first 10–50 ns, rarely benefit from extended sampling, and short simulations are sufficient for validation or minor relaxation. By contrast, models of intermediate quality can be triaged based on their early non-canonical fidelity trajectories: a rapid upward trend signals refinement potential and justifies longer runs, whereas an early decline strongly predicts failure and suggests termination. Poor or topologically flawed starting structures, typically with very high RMSD or collapsed non-canonical networks, show no meaningful recovery in this early window and should not be pursued with MD at all. Finally, catastrophic early collapse is an unmistakable diagnostic for immediate termination. These decision points make it possible to minimize computational cost while preserving the refinement potential of MD, ensuring that resources are focused only on models with a realistic chance of improvement.

How to recognize model quality in practice—even without a native reference—can be guided by indirect indicators. Consensus among multiple prediction methods, agreement with reliable secondary structure predictions (e.g., SHAPE probing or covariation analysis), or consistency with experimental restraints (e.g., cryo-EM density, NMR NOEs) are strong signs of reliability. During equilibration and in the first tens of nanoseconds of MD, stability of canonical base pairs (INF_wc close to 1.0 relative to t=0), a flat RMSD trajectory, and the absence of rapid collapse in stacking or non-canonical contacts are additional indicators of a robust starting model. Conversely, large RMSD jumps, early loss of canonical pairs, or disordered loop collapse signal that the model is not suitable for refinement.

General decision pathways are summarized in Table 1, while a detailed diagnostic checklist is provided in Table 2. Together, these resources provide a practical framework for when and how to apply MD simulations in RNA structure modeling: short runs for validation of good models, selective extension when early improvement is observed, and immediate termination when instability is evident.

## Conclusions

This study demonstrates that MD simulations are not a universally reliable refinement strategy for RNA structural models and are best applied selectively. The success of MD is critically dependent on the initial quality of the starting model, often more so than the target’s difficulty classification.

While short MD simulations (10–50 ns) can provide modest improvements in interaction network fidelity for well-predicted RNA structures, they were generally unable to correct major errors in poorly modeled structures. Extended simulations (>100 ns) tended to increase variability, disrupted non-Watson–Crick interactions, and often degraded model accuracy. High-quality starting models—even for’Difficult’ or’Non-natural’ targets—could remain remarkably stable or undergo partial refinement, whereas poor models of’Easy’ showed limited benefit.

Given these findings, and considering the substantial computational cost, MD is best reserved as a precision tool for fine-tuning or validating high-quality models. Its use should be carefully controlled through pre-selection of starting structures, optimal simulation timescales, and early-phase diagnostics such as INF_nwc monitoring. In cases of low-quality models, alternative strategies (hybrid modeling, experimental restraints, fragment-based approaches) should be prioritized.

Ultimately, MD is most powerful when deployed selectively: as a diagnostic and refinement tool for models already close to the native state, not as a universal solution. Based on our CASP15 analysis, we provide practical guidelines for incorporating short MD simulations into RNA modeling pipelines—highlighting when they are useful, when they offer limited benefit, and how to apply them in a cost-effective manner. Future improvements in force fields, integration of experimental data, and hybrid modeling approaches may expand the utility of MD, but at present, its application should remain selective, evidence-based, and mindful of computational cost.

## Acknowledgements

S.K., C.N. and S.P acknowledge funding from the National Science Centre, Poland (SHENG 2021/40/Q/NZ2/00078). We gratefully acknowledge Polish high-performance computing infrastructure PLGrid (HPC Center: ACK Cyfronet AGH) for providing computer facilities and support within computational grant no. PLG/2025/017952.

## Data Availability

All molecular dynamics simulation snapshots generated in this study are available at Zenodo under accession <u>10.5281/zenodo.16915355</u>. Reference/native and starting CASP15 RNA models are available from the official CASP website: https://predictioncenter.org/casp15/

## Declaration of generative AI and AI-assisted technologies in the writing process

During the preparation of this work the authors used ChatGPT, Google AI studio and Claude to proof-read the text to enhance the quality of the language. After using these tools, the authors reviewed and edited the content as needed and take full responsibility for the content of the publication.

## Supplementary Data

Supplementary Table S1: CASP15 RNA Models for Analysis

Supplementary Figure S1: Time series plots of Interaction Network Fidelity (INF) scores over the 300 ns simulation

Supplementary Figure S2: Dynamics of Interaction Network Fidelity.

Supplementary Figure S3: Quantitative Summary of Interaction Network Fidelity Changes Over Time.

Supplementary Figure S4: Conformational landscape for R1117 simulations projected onto the dominant tICA components.

Supplementary Figure S5: Conformational landscape for R1107 simulations projected onto the dominant tICA components.

Supplementary Figure S6: Computational cost of RNA molecular dynamics simulations.

Supplementary Figure S7: Landscape of ‘Difficult’ target R1116 (Cloverleaf RNA). Supplementary Figure S8: Landscape of ‘Non-natural’ target R1128 (Paranemic crossover triangle).

### Supplementary Case Studies

Difficult target with transient nWC improvement followed by degradation Non-natural target retaining secondary structure but losing tertiary contacts

## Notes

https://doi.org/10.5281/zenodo.16915355

### Competing Interest Statement

The authors have declared no competing interest.

