## Supplementary for "When Does Molecular Dynamics Improve RNA Models? Insights from CASP15 and Practical Guidelines"

**Supplementary Table S1: CASP15 RNA Models for Analysis**

| Target | Length (nt) | Difficulty | Group_ Model# | RMSD (Å) |
|---|---|---|---|---|
| R1107 | 69 | Medium | TS232_1 | 4.52 |
| | | | TS287_2 | 6.48 |
| | | | TS081_4 | 8.80 |
| | | | TS128_3 | 5.98 |
| | | | TS416_5 | 6.51 |
| | | | TS392_1 | 16.65 |
| R1108 | 69 | Medium | TS232_4 | 4.49 |
| | | | TS287_2 | 5.95 |
| | | | TS081_4 | 8.48 |
| | | | TS128_3 | 5.48 |
| | | | TS416_3 | 4.80 |
| R1116 | 157 | Difficult | TS232_5 | 17.26 |
| | | | TS287_4 | 17.99 |
| | | | TS081_1 | 12.65 |
| | | | TS416_3 | 12.69 |
| | | | TS392_1 | 21.22 |
| R1117 | 30 | Easy | TS232_1 | 2.27 |
| | | | TS287_4 | 2.01 |
| | | | TS081_1 | 2.74 |
| | | | TS128_1 | 2.43 |
| | | | TS416_1 | 2.88 |
| | | | TS392_5 | 4.70 |
| R1126 | 363 | Non-natural | TS232_4 | 8.76 |
| | | | TS287_2 | 12.64 |
| | | | TS081_1 | 19.95 |
| | | | TS128_1 | 33.42 |
| | | | TS416_2 | 41.93 |
| | | | TS392_1 | 41.12 |
| R1128 | 238 | Non-natural | TS232_1 | 4.33 |
| | | | TS287_1 | 6.68 |
| | | | TS081_5 | 14.61 |
| | | | TS128_3 | 33.02 |
| | | | TS416_2 | 14.27 |
| | | | TS392_5 | 21.07 |
| R1136 | 374 | Non-natural | TS232_3 | 7.25 |
| | | | TS287_4 | 10.94 |
| | | | TS081_2 | 10.97 |
| | | | TS128_1 | 13.23 |
| | | | TS416_2 | 32.25 |

| | | | | |
|---|---|---|---|---|
| | | | TS392_2 | 39.23 |
| R1149 | 124 | Difficult | TS232_5 | 10.46 |
| | | | TS287_4 | 14.01 |
| | | | TS081_2 | 18.18 |
| | | | TS128_1 | 7.36 |
| | | | TS416_3 | 7.66 |
| | | | TS392_4 | 12.70 |
| R1156 | 135 | Difficult | TS232_3 | 7.61 |
| | | | TS287_1 | 11.02 |
| | | | TS081_4 | 17.06 |
| | | | TS128_5 | 5.37 |
| | | | TS416_1 | 17.65 |
| | | | TS392_5 | 14.81 |

[#] The models submitted by the groups Alchemy_RNA2 (TS232), Chen (TS287), RNApolis (TS081), Genesilico (TS128), Alchemy_RNA (TS416), and LCBio (TS392) were used in this study.

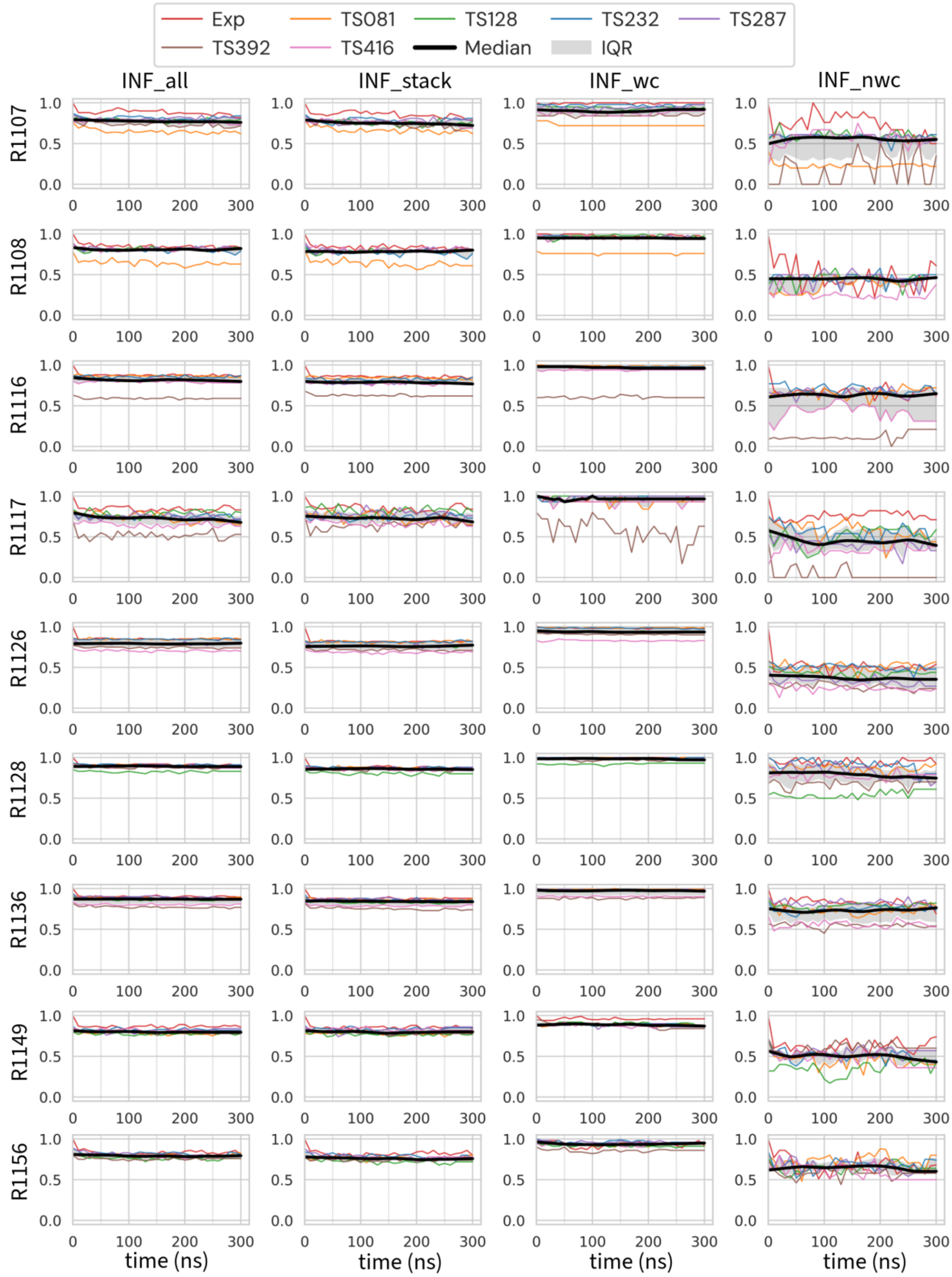


**Supplementary Figure S1: Time series plots of Interaction Network Fidelity (INF) scores over the 300 ns simulation.** The four subplots show the overall fidelity (inf_all), stacking (inf_stack), Watson-Crick (inf_wc), and non-Watson-Crick (inf_nwc) components. Individual colored lines represent each simulation, corresponding to the arrows in panel A. The solid black line indicates the median INF score of all submitted

CASP models (excluding the experimental reference). The shaded grey area represents the interquartile range (IQR), which illustrates the behavior of the central 50% of the models by spanning the 25th to 75th percentiles.

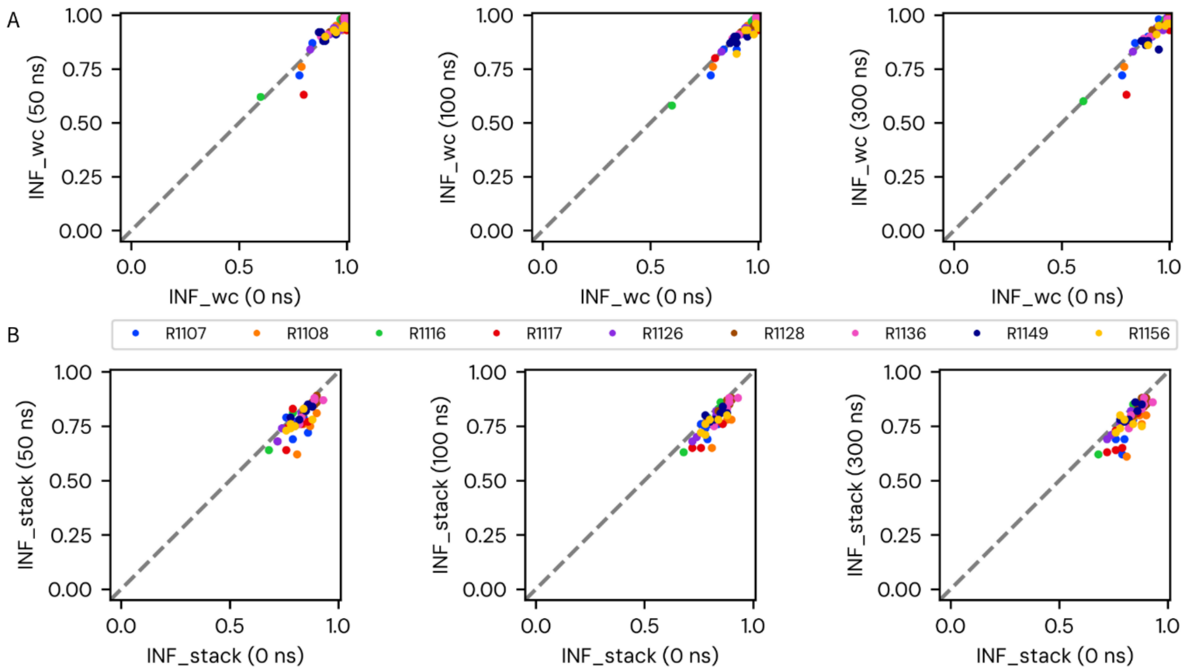


**Supplementary Figure S2: Dynamics of Interaction Network Fidelity.** (A) Scatter plots comparing the canonical interaction fidelity (inf_wc) of each simulated model at three time points (50, 100, and 300 ns) against its initial fidelity (0 ns). Each point represents a single model, color-coded by its RNA target. The dashed diagonal line represents perfect stability; points below the line indicate degradation. (B) Scatter plots for the stacking interaction fidelity (inf_stack), showing the same comparison as in (A). The significant scatter on both sides of the identity line highlights the dynamic nature of the non-canonical network, which is the locus of both refinement (points above the line) and degradation (points below). The inf_all and inf_nwc components are presented in Figure 2.

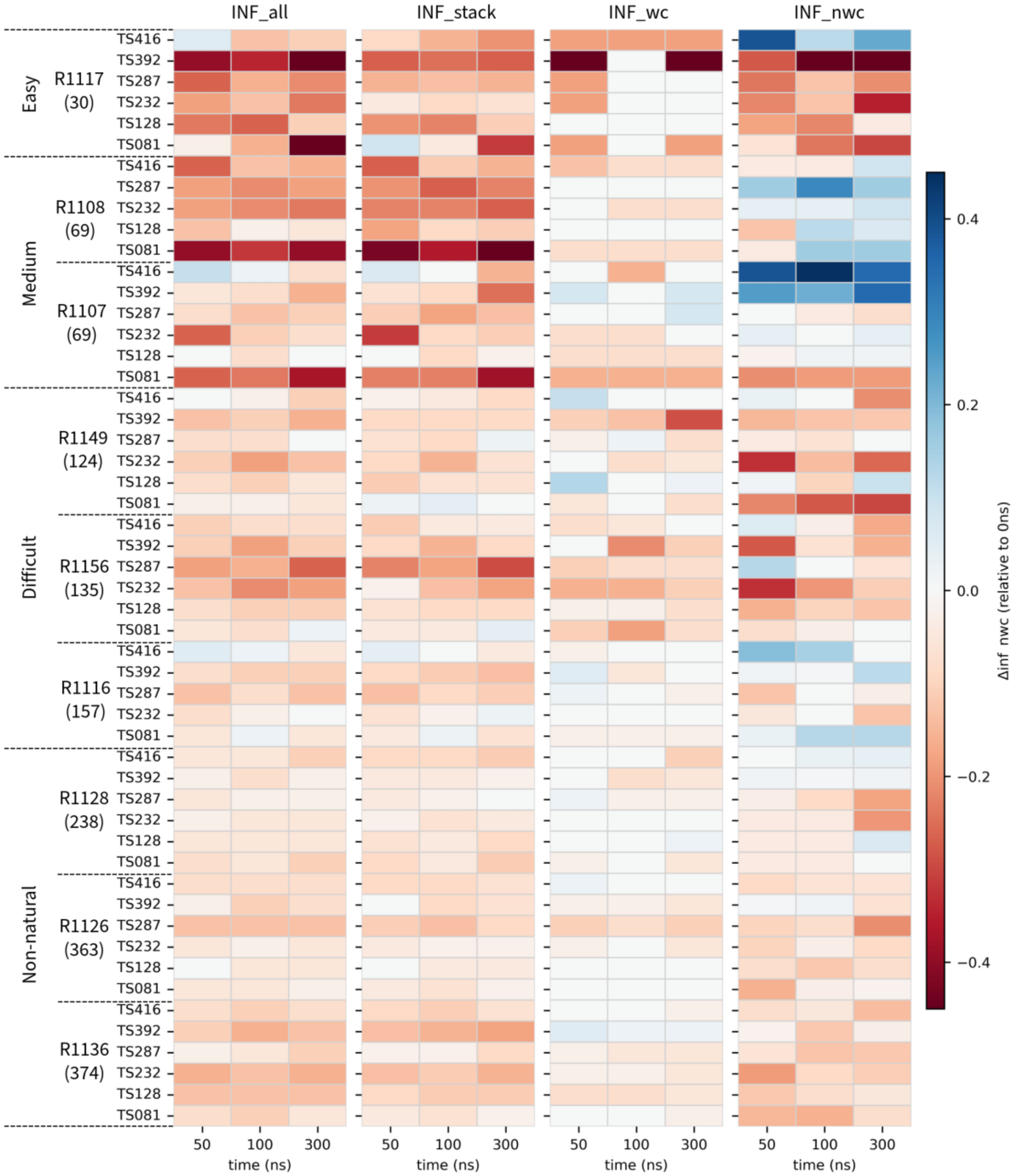


**Supplementary Figure S3: Quantitative Summary of Interaction Network Fidelity Changes Over Time.** The figure presents a quantitative heatmap of the change (ΔINF) in Interaction Network Fidelity for each simulated model at three time points (50, 100, and 300 ns) relative to its initial state (0 ns). Each row corresponds to a single model, grouped by RNA target and its difficulty classification. The four main columns represent the different INF components (inf_all, inf_stack, inf_wc, and inf_nwc). The color of each cell corresponds to the numerical value of the change in INF (ΔINF), as indicated by the color bar. Blue shades represent an increase in fidelity, while red shades represent a decrease. This quantitative view complements the categorical summary in the main text, providing the precise magnitude of the observed refinement or degradation for each model's interaction network.

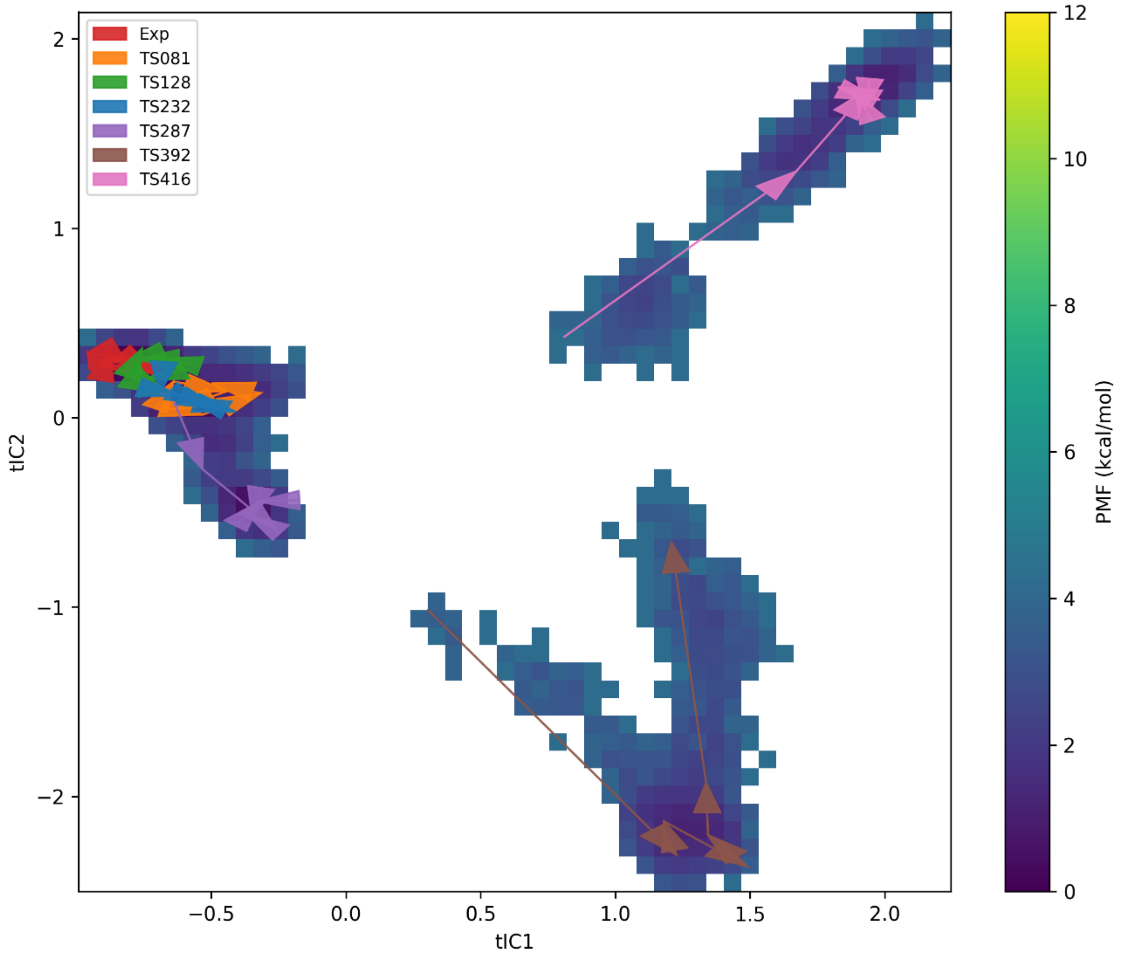


**Supplementary Figure S4: Conformational landscape for R1117 simulations projected onto the dominant tICA components.** Combined 2D Potential of Mean Force (PMF) landscape for all R1117 simulations, projected onto the first (tIC1) and second (tIC2) time-lagged independent components. The landscape energy is shown in kcal/mol, with low-energy basins in purple and high-energy barriers in yellow. Arrows trace the conformational pathway for the native reference (Exp) and each CASP model from the start (tail) to the end (head) of the 300 ns simulation. This figure shows the dynamics along the two most dominant collective motions of the system, complementing the higher-order tIC projection shown in the main text (Figure 4).

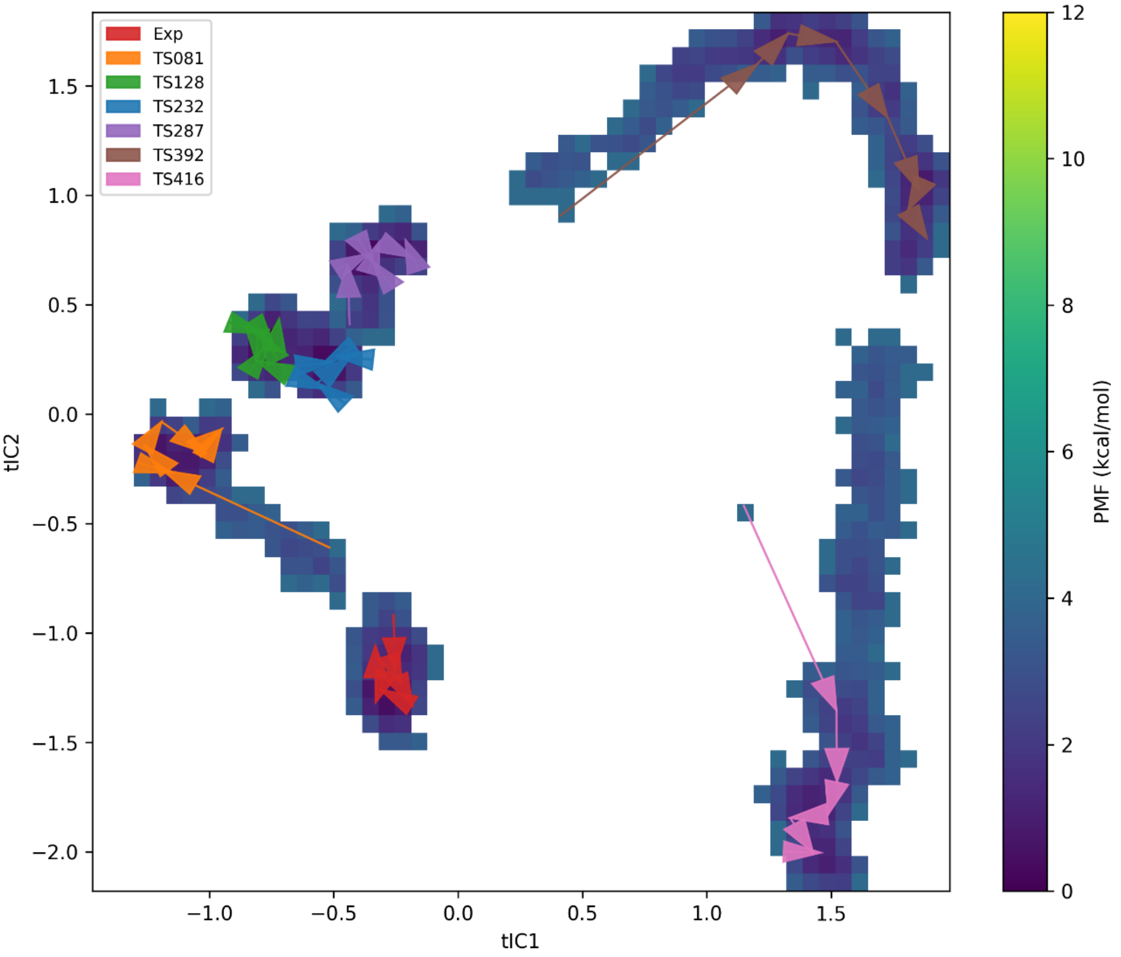


**Supplementary Figure S5: Conformational landscape for R1107 simulations projected onto the dominant tICA components.** Combined 2D Potential of Mean Force (PMF) landscape for all R1107 simulations, projected onto the first (tIC1) and second (tIC2) time-lagged independent components. The landscape energy is shown in kcal/mol, with low-energy basins in purple and high-energy barriers in yellow. Arrows trace the conformational pathway for the native reference (Exp) and each CASP model from the start (tail) to the end (head) of the 300 ns simulation. This figure shows the dynamics along the two most dominant collective motions of the system, complementing the higher-order tIC projection shown in the main text (Figure 5).

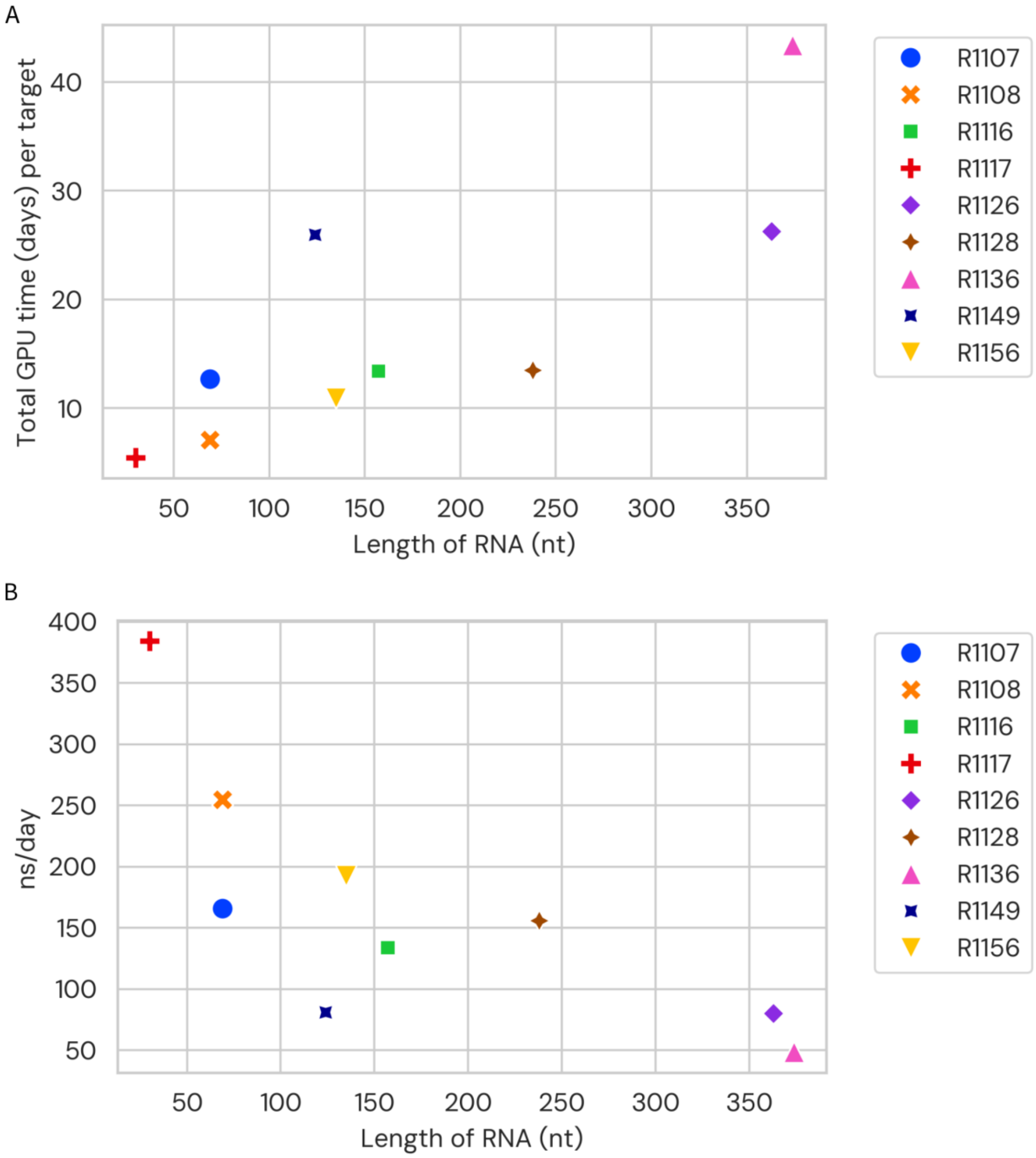


**Supplementary Figure S6: Computational cost of RNA molecular dynamics simulations.** (A) The total aggregate GPU time required for each target with seven models, with the exception of six each for R1108 and R1116. The total computational time per target ranged from approximately 5.5 GPU-days for the smallest target (R1117) to 43.4 GPU-days for the largest (R1136). The simulations for all 61 models required a total of 158.6 GPU-days. (B) The simulation performance for each target, measured in nanoseconds (ns) of simulation generated per day. This plot shows a strong inverse correlation between RNA length and simulation speed; the smallest systems run the fastest (e.g., R1117 at 384.5 ns/day), while the largest systems are the slowest (e.g., R1136 at 48.4 ns/day). All simulations were performed on an NVIDIA GH200 (96 GB) GPU.

**Supplementary Case Studies**

**Difficult target with transient nWC improvement followed by degradation**

Target R1116, a 'Difficult' Cloverleaf RNA, displayed brief and partial recovery of non-canonical contacts in one model during the first half of MD, but these gains were lost in the second half, resulting in a net degradation of interaction fidelity. The starting models had an RMSD in the range of 12.65 – 21.22 Å. To explore conformational dynamics and refinement behavior, we projected all simulations onto a shared 2D potential of mean force (PMF) landscape (Supplementary Figure S7) and analyzed interaction fidelity over time (Supplementary Figure S1).

Overall, the simulations revealed a consistent trend of degradation or stagnation. Even the highest-scoring models struggled to maintain native-like contacts, especially in the non-canonical interaction network. For example, TS232 (blue arrow) and TS287 (purple arrow) both began with high overall fidelity (INF_all ≈ 0.87) but saw their INF_nwc drop from 0.77 to 0.64 and 0.74 to 0.61, respectively (Supplementary Figure S1). This highlights a key limitation of MD-based refinement for difficult targets: instead of repairing weak regions, simulations often erode delicate tertiary interactions.

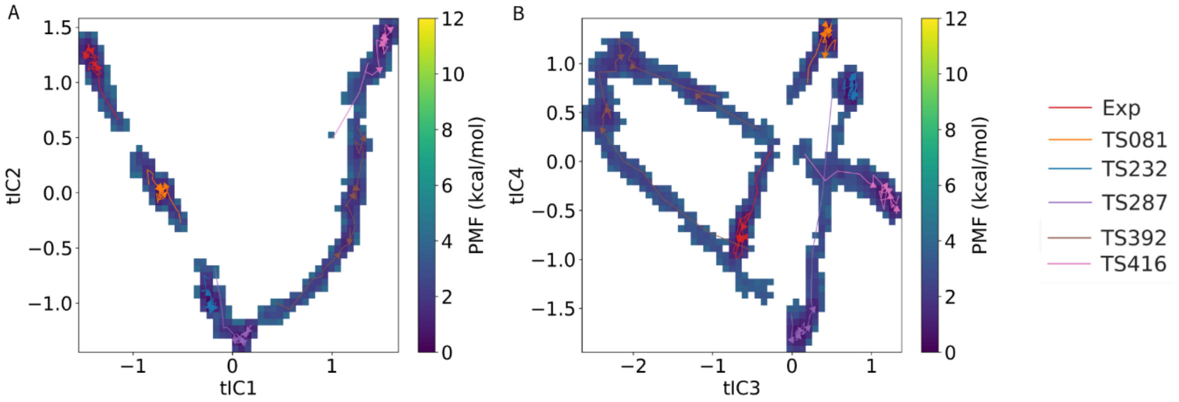


**Supplementary Figure S7: Landscape of 'Difficult' target R1116 (Cloverleaf RNA).** Combined 2D Potential of Mean Force (PMF) landscape for all MD simulations of R1116, projected onto (A) tIC1 and tIC2 and (B) tIC3 and tIC4. Free energy is shown in kcal/mol, with low-energy basins in purple and high-energy regions in yellow. Arrows trace model trajectories from 0 to 300 ns, with the experimental reference in red.

More severely flawed models showed distinct failure modes. TS392 (brown arrow) represents a case of complete collapse. It started with low overall fidelity (INF_all = 0.63) and a broken Watson–Crick pairing network (INF_wc = 0.60). As shown in Supplementary Figure S4A, the model remained trapped in a non-native basin for the entire 300 ns, and failed to recover any meaningful interactions (Supplementary Figure S1).

TS416 (pink arrow) illustrates a more complex trajectory. It began with moderate global fidelity (INF_all = 0.80) but weak non-canonical contacts (INF_nwc = 0.31). During the simulation, INF_nwc rose steadily to 0.58 by ~120 ns, suggesting transient improvement. However, these gains proved unstable: the score declined sharply in the second half of the trajectory, ending near its starting value. This case underscores a common outcome in difficult targets—MD can occasionally sample better conformations, but converting them into stable, refined structures remains rare and unpredictable.

### Non-natural target retaining secondary structure but losing tertiary contacts

Target R1128, a designed RNA origami structure, maintained nearly perfect Watson–Crick pairing and stacking throughout MD, yet consistently lost key non-canonical contacts that are essential for stabilizing its tertiary fold. As a 'Non-natural' target, R1128 poses a distinct challenge: it requires de novo prediction methods to solve an entirely novel topology without relying on evolutionary constraints. CASP15 results showed that the best models for such targets often came from human-guided groups, however, with a wide range of 4.33 - 21.07 Å.

The PMF landscape (Supplementary Figure S8) shows a clear separation between the experimental reference and the predicted models. The reference simulation (red arrow) remained confined to a well-defined low-energy basin, indicating exceptional structural stability. In contrast, all predicted models sampled distinct and less stable regions, failing to converge on the native basin.

This divergence is mirrored in the interaction fidelity time series (Supplementary Figure S1). Watson–Crick pairing (INF_wc) and stacking (INF_stack) were consistently strong across models. For instance, TS232 (blue arrow) and TS081 (orange) maintained perfect Watson–Crick networks (INF_wc = 1.0) throughout the simulation. These results confirm that predictors successfully captured the secondary structure.

However, the non-canonical interaction network (INF_nwc) told a different story. All models started with lower inf_nwc than the experimental reference, and most declined further during simulation. TS232 began with INF_nwc = 1.0 but dropped to 0.80. TS081 fell from 0.92 to 0.89, and TS287 (purple) dropped from 0.88 to 0.70. While these drops may seem moderate, they reveal an inability to preserve the specific non-canonical architecture that stabilizes the native fold.

The case of R1128 underscores a central limitation in current RNA structure prediction pipelines. Accurate secondary structure is necessary but not sufficient. For complex, designed RNAs, the critical determinant of correctness lies in the non-canonical tertiary contacts—interactions that remain elusive to both predictors and refinement via molecular dynamics.

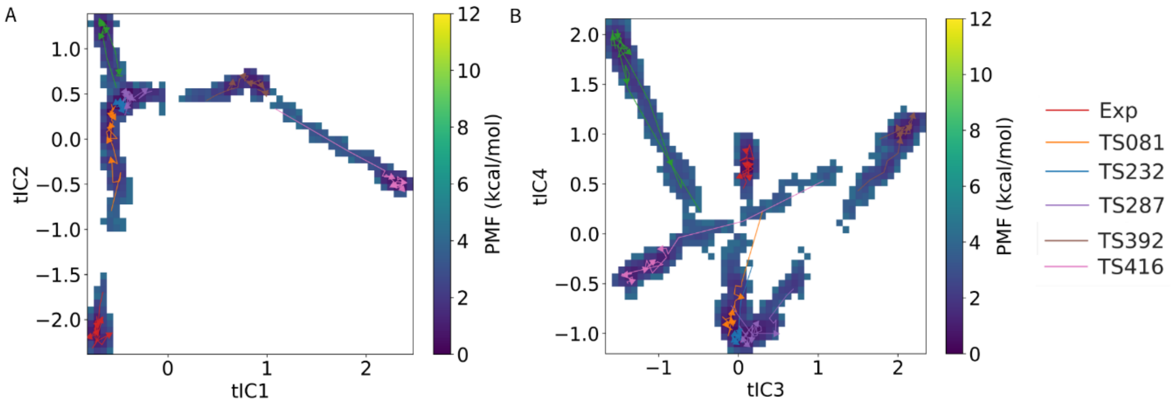

**Supplementary Figure S8: Landscape of 'Non-natural' target R1128 (Paranemic crossover triangle).** Combined 2D Potential of Mean Force (PMF) landscape for all MD simulations of R1128, projected onto (A) tIC1 and tIC2 and (B) tIC3 and tIC4. Free energy is shown in kcal/mol, with low-energy basins in purple and high-energy regions in yellow. Arrows trace conformational pathways of the experimental structure (red) and CASP models from 0 to 300 ns.
